# Actomyosin Contractility Drives Bile Regurgitation as an Early Homeostatic Response to Increased Biliary Pressure in Obstructive Cholestasis

**DOI:** 10.1101/077792

**Authors:** Kapish Gupta, Qiushi Li, Jun Jun Fan, Eliza Li Shan Fong, Ziwei Song, Shupei Mo, Haoyu Tang, Inn Chuan Ng, Chan Way Ng, Pornteera Pawijit, Shuangmu Zhuo, Chen-Yuan Dong, Boon Chuan Low, Aileen Wee, Yock Young Dan, Pakorn Kanchanawong, Peter So, Virgile Viasnoff, Hanry Yu

**Affiliations:** Mechanobiology Institute, National University of Singapore, Singapore; National University of Singapore Research Institute, Singapore; Institute of Bioengineering and Nanotechnology, Agency for Science, Technology and Research (A*STAR), Singapore; BioSyM, Singapore-MIT Alliance for Research and Technology, Singapore; Department of Orthopaedics, Xijing Hospital, Fourth Military Medical University, China; Department of Physiology, National University of Singapore, Singapore; Department of Biomedical Engineering, National University of Singapore, Singapore; NUS Graduate School of Integrative Sciences and Engineering, National University of Singapore, Singapore 117456; Fujian Normal University, Fuzhou, Fujian, China; Department of Physics, National Taiwan University, Taiwan; Department of Biological Sciences, National University of Singapore; Department of Pathology, National University of Singapore, Singapore; Division of Gastroenterology and Hepatology, National University Hospital, Singapore; CNRS UMI3639, 5A Engineering Drive 1 117411 Singapore; Department of Gastroenterology, Nanfang Hospital, Southern Medical University, Guangzhou, China

**Keywords:** blebs, vesicles, actomyosin cortex, hepatocytes, bile canaliculi

## Abstract

A wide range of liver diseases manifest as biliary obstruction, or cholestasis. However, the sequence of molecular events triggered as part of the early hepatocellular homeostatic response to abnormal elevations in biliary pressure remains poorly elucidated. Bile canaliculi are dynamic luminal structures that undergo actomyosin-mediated periodic contractions to propel secreted bile. Additionally, pericanalicular actin is accumulated during obstructive cholestasis. Therefore, we hypothesize that the pericanalicular actin cortex undergoes significant remodeling as a regulatory response against increased biliary pressure. Here, we report that, actomyosin contractility induces transient deformations along the canalicular membrane, a process we have termed inward blebbing. We show that these membrane intrusions are initiated by local ruptures in the pericanalicular actin cortex, and they typically retract following repair by actin polymerization and actomyosin contraction. However, above a certain osmotic pressure threshold, these inward blebs pinch away from the canalicular membrane into the hepatocyte cytoplasm as large vesicles (2-8 µm). Importantly, we show that these vesicles aid in the regurgitation of bile from the canalicular system. Conclusion: Actomyosin contractility induces the formation of bile-regurgitative vesicles, thus serving as an early homeostatic mechanism against increased biliary pressure during cholestasis.

## Introduction

The biliary function of the liver is critical for survival, serving to eliminate toxic endo- and xenobiotics, cholesterol, and inflammatory mediators.[1] The apical membranes of adjacent hepatocytes form the bile canalicular lumen, an intercellular structure surrounded by a dynamic pericanalicular actin cortex (PAC), which actively contracts to propel secreted biliary fluid towards the bile ducts.[2, 3] A variety of liver diseases result in impaired bile flow, or obstructive cholestasis.[4–7] These include extrahepatic etiologies such as biliary strictures, stones and biliary atresia in infants; as well as intrahepatic causes that include primary biliary cirrhosis, vanishing duct syndrome and, alcoholic and viral hepatitis. Bile stasis and backpressure increases liver and serum bile acid levels that results in liver toxicity and fibrosis, which may eventually progress to decompensated cirrhosis, mandating the need for a liver transplant. [7]

Several pathological changes that are effected consequent to increased biliary pressure have been identified, including changes in transporter expression which reduce the uptake and increase the basolateral export of bile acids,[6] and the accumulation of actin along the canalicular membrane. [8, 9] Indeed, the thickening of the PAC observed in common bile duct ligation rodent models has also been observed in patients with biliary atresia.[10] However, these reports have largely been correlative; little is known about the exact sequence of events that is triggered to effect homeostatic regulation of abnormal elevations in biliary pressure.

More than 40 years ago, Matter and colleagues reported the presence of vacuoles containing horseradish peroxidase (HRP) in the hepatocyte cytoplasm upon retrograde injection of HRP through the common bile duct. [11] Though not proven, these results led to a proposed model whereby these vacuoles were part of a diacytotic-based process of bile regurgitation that enable the transport of bile from the canalicular to the sinusoidal surface during increased biliary pressure. In another study, Watanabe and colleagues monitored the process of bile regurgitation from the canalicular space through the hepatocyte cytoplasm into the sinusoids following common bile duct ligation (BDL), thus establishing the transcellular pathway as the main homeostatic mechanism that protects hepatocytes from bile toxicity during increased biliary pressure. [12] Together, these two studies implicate the transcellular transport of bile as an immediate homeostatic mechanism triggered by increased biliary pressure. However, the precise molecular machinery and sequential events underlying this phenomenon is still poorly understood.

Approaching obstructive cholestasis as a disease of aberrant cellular mechanics, the known motility of the canalicular network and reported involvement of the PAC in bile flow and bile stasis suggest that cytoskeletal changes in the PAC may be involved as an early homeostatic mechanism to counteract elevations in biliary pressure; the failure of which then results in further adaptive changes. [3] In this study, we detailed in real-time the sequence of events that occur immediately following induced elevations in biliary pressure, and demonstrated the role of actomyosin contractility in facilitating bile regurgitation. A mechanistic understanding of the early homeostatic response triggered to relieve intracanalicular pressure (ICP) may lead to the identification of therapeutic targets to prevent or treat obstructive cholestasis.

## Materials and Methods

### Maintenance and *in vivo* imaging of LifeAct mice

All animal experiments were approved and in accordance to the guidelines by the Institutional Animal Care and Use Committee (IACUC) of the Agency for Science, Technology and Research (A*STAR) in Singapore. Transgenic LifeAct-GFP mice[13] (20 weeks old, average body weight of 28 g) were anesthetized by intraperitoneal injection of ketamine (100 mg/kg) and xylazine (10 mg/kg). An intravital imaging window chamber was mounted on the abdomen as previously described.[14] Intravital imaging was performed with a titanium-sapphire laser (Tsunami, SpectraPhysics, Mountain View, California) with a 488 nm output. The laser was scanned using a x-y mirror scanning system (Model 6220, Cambridge Technology, Cambridge, Massachusetts) and guided towards the modified inverted microscope. A high power objective (Plan Fluor ELWD water 40X, NA 0.45, Nikon) was used. After passing through the primary dichroic mirror, the GFP fluorescence signal was detected with a 488 nm bandwidth using additional band pass filters (HQ590/80, ChromaTechnology). Each optical scan is composed of 512 by 512 pixels and took approximately 1s to complete.

### Bile duct ligation

Mice were anesthetized by intraperitoneal injection of ketamine (100 mg/kg) and xylazine (10 mg/kg). The common bile duct was ligated using double surgical knots below the bifurcation and one single knot above the pancreas. Following ligation, the window chamber was attached to the animal and imaging was performed 1 h after the procedure as described above. Three mice from each of the control and bile duct-ligated groups were imaged in this study.

### Isolation and sandwich culture of hepatocytes

Hepatocytes were isolated from male Wistar rats using a previously described two-step *in situ* collagenase perfusion method. [15] Isolated hepatocytes were cultured in collagen sandwich configuration. Hepatocytes were either transfected with various florescent tagged proteins or were exposed to different concentration of blebbistatin, cytochalasin D or ursodeoxycholic acid (UDCA) to alter canalicular dynamics. For investigations of the role of BCV, hepatocytes were exposed to cholyl-lysyl-fluorescein (CLF). Details of hepatocyte isolation, culture and treatment of hepatocytes are discussed in Supporting Information.

## Results

### Observation of inward bleb formation in normal BC *in vivo*

Using intravital microscopy (Figure 1 A-C) to image the LifeAct probe of filamentous (f)-actin in the mouse liver (Figure 1–4), we imaged the liver surface (2-3 cell layers in depth). Besides the epithelium and sinusoidal network, we were also able to identify the BC network which had a greater intensity as actin is highly expressed at the apical surface of hepatocytes [16]. As the actin in the BC network gave a much stronger signal than the surrounding tissue, we segmented the BC using simple threshold (Supporting Figure 1). By visual inspection, we observed that the average diameter of BC in the control group was 1.48±0.14 µm (n=20), excluding the possibility that these structures were bile ducts, which are known to have diameters greater than 10 µm (Figure 1 D and E). [17–19] Based on our knowledge, this study demonstrates for the first time the ability to live image the BC network with such resolution. With the ability to study the *in vivo* BC network in real-time, we first sought to characterize the periodic cycles of bile canalicular expansion and contraction. In agreement with previous literature,[3, 20–22] BC were observed to be highly dynamic structures. However, beyond the previously described global motility of BC, we discovered that the bile canalicular surface is also remarkably dynamic on the local scale. During the BC expansion phase, we observed the formation of membrane herniations (1-2 µm in diameter with lifetime of approximately 2 min) which occur randomly and dynamically on the bile canalicular surface. These bleb-like structures intrude inwardly into the hepatocyte cytoplasm and retract back towards the bile canalicular lumen (Figure 2C and 1). The morphology and dimensions of these dynamic structures are reminiscent of classical membrane blebs, which are outward membrane protrusions arising from local cortical weakening and a temporary decoupling of the actin cortex from the plasma membrane.[23–25] However, in marked contrast to classical extruding blebs, these membrane intrusions exhibit an opposite directionality. Therefore, we termed these dynamic structures, inward blebs (IB).

**Figure 1:**
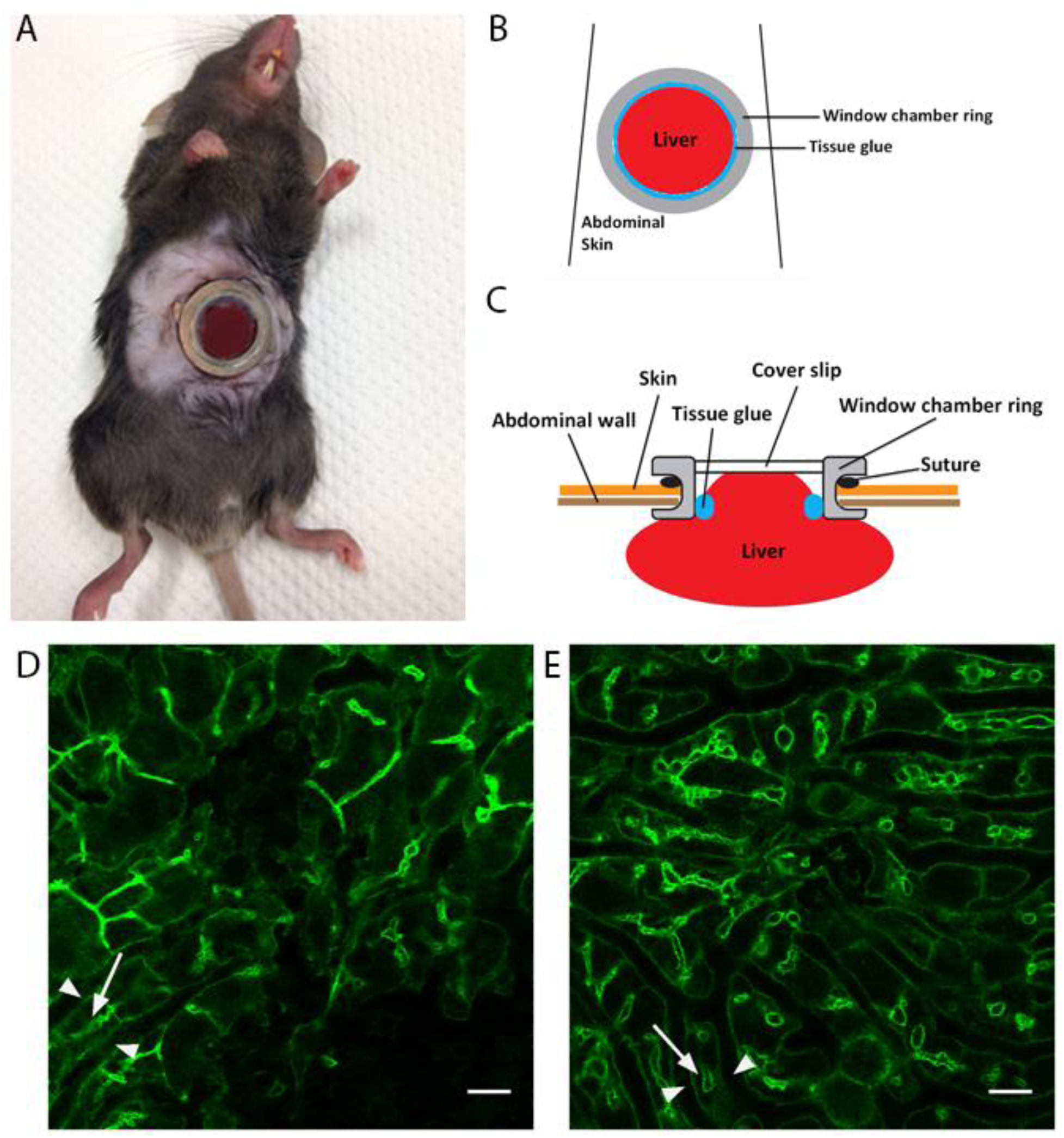
*In vivo* imaging of mouse liver (F-actin is labeled green) using window chamber reveals intricate bile canaliculi network in normal and bile duct-ligated LifeAct-GFP mice. Confocal imaging was performed using the intravital imaging set up as shown in (A) to (C). (A) A mouse with coverslip glued to the imaging setup ready for imaging. (B) Schematic of set up showing window chamber over liver. (C) Side view schematic of set-up. A coverslip is placed on the exposed mouse liver during imaging. Using this set up, we were able to view 2-3 layers of hepatocytes below the liver surface. (D) shows a confocal slice of the liver in normal GFP-LifeAct mice and (E) shows a confocal slice of the liver in bile duct-ligated GFP-LifeAct mice. Bile canaliculi (white arrow) were identified as actin structures that formed between hepatocytes. As can be seen in (D) and (E), actin localized along the hepatocyte cell cortex (white arrowhead) and bile canaliculi (white arrow), but much more actin was localized along the bile canaliculi as expected. Therefore, the bile canalicular structures appear as distinct structures when imaged. Also, bile canaliculi in bile duct-ligated mice were more dilated compared to normal mice. In normal mice, the average diameter of bile canaliculi was 1.48±0.14 µm (n=20) and that in bile duct-ligated mice was 3.55±0.3 µm (n=20).

**Figure 2:**
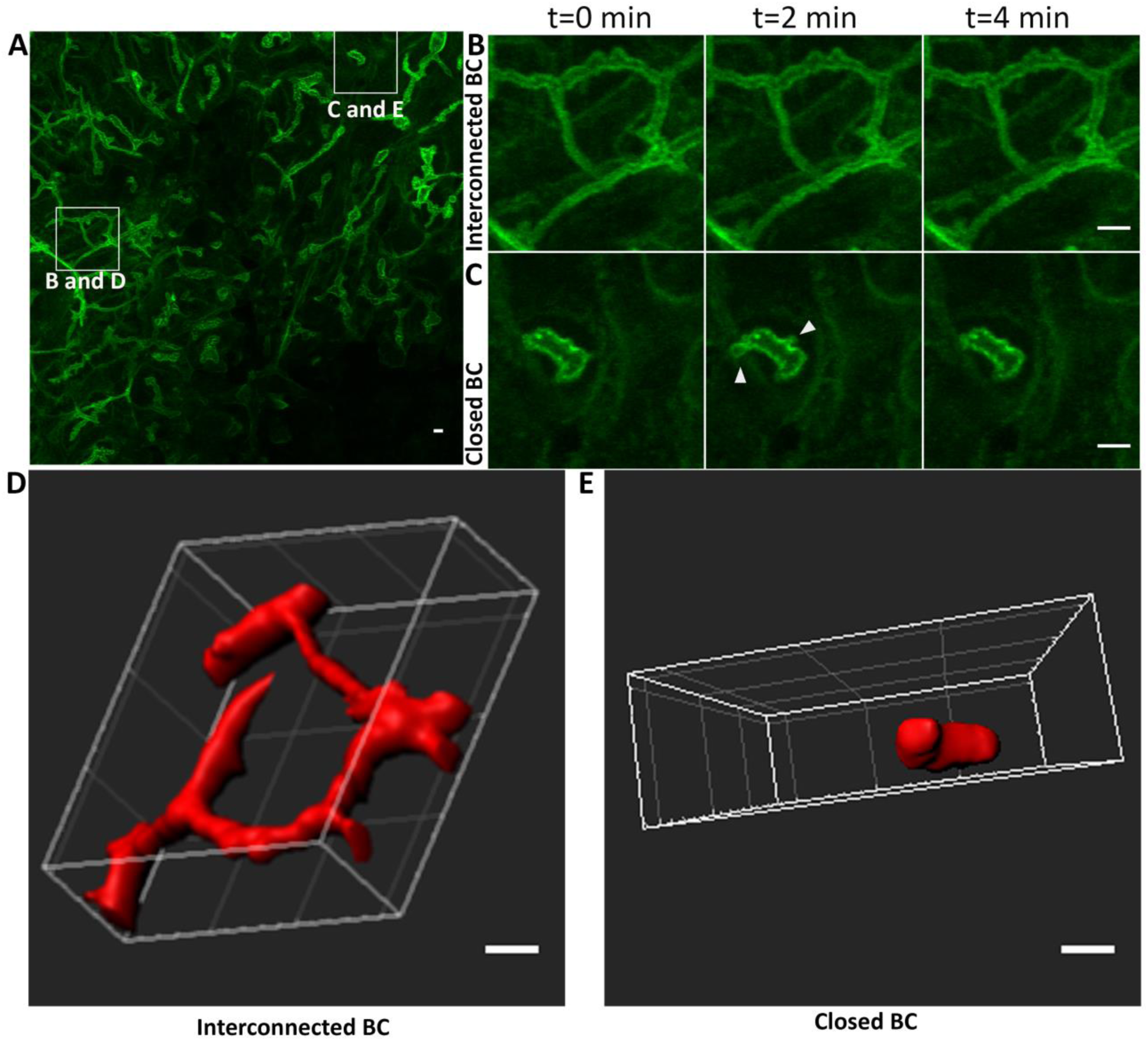
Inward blebbing is observed only at closed or isolated bile canaliculi. (A) shows the Z-projection of confocal slices taken of the liver in normal GFP-LifeAct mice (a representative slice in the stack is shown in Figure 1E; additional slices are shown in Supporting Information). Note that in the Z-projection, only the bile canaliculi are distinctly visible as the amount of actin is much higher along the bile canalicular surface as compared to other sections of the hepatocyte cortex. Inward blebs were not observed along long and interconnected bile canaliculi as magnified in (B). Instead, inward blebs were only found at the terminal ends of the bile canalicular network, or along short and isolated bile canaliculi, magnified in (C). (B) and (C) show time lapse images (at 2 min intervals) of the regions highlighted by the squares in (A). White arrowheads in (C) indicate the formation of inward blebs. (D) and (E) are representative reconstructed 3D images of interconnected (B) and closed (C) bile canaliculi. Supporting Figures 2 and 3 illustrate how interconnected and closed bile canaliculi were determined. Bile canaliculi were considered closed only if they were not connected to other canaliculi when viewed after 3D reconstruction of confocal slices. All scale bars = 5 µm.

### Biliary pressure promotes IB vesicularization *in vivo*

As classical extruding blebs generally arise due to an increase in intracellular hydrostatic pressure, [23, 26, 27] we asked if IB occurred at regions of increased canalicular pressure in the BC network. To investigate this, we examined fluorescence micrographs of mouse liver expressing LifeAct (Figure 2). We found that the frequency of blebbing at the terminal ends of the BC network or, along the short and isolated BC, was 2.5-fold higher than the global blebbing frequency (Figure 3D) in the control mice. We defined global blebbing frequency as the total blebbing frequency at both interconnected (Figure 2B, 2D and Supporting Figure 2) and closed (Figure 2C, 2E and Supporting Figure 3) BC. We categorized BC as either closed or interconnected based on 3D reconstruction of confocal z stacks, as shown in Figure 2D and E and Supporting Figures 2,3 and 4. This result suggests that IB preferentially occurs at the terminal ends of the BC network or, along short and isolated BC where canalicular pressure is expected to be higher, rather than in the middle of long and interconnected BC (1). Importantly, when biliary pressure was increased by ligating the common bile duct (Figure 3), [28] not only did the average BC diameter increase (3.55±0.3 µm, n=20), we observed an increase in the frequency of IB and found that the global blebbing frequency in BDL mice was similar to the blebbing frequency at the canalicular ends in normal mice (Figure 3D). Increase in BC diameter in BDL is in accordance with earlier published reports [29]. Furthermore, in BDL mice, IB were no longer confined to regions where BC were short and isolated (Figure 3C and 4B), but were also found along the long and interconnected BC (Figure 3B and Supporting Figure 4), indicating that IB formation is indeed consequent to increased biliary pressure (Figure 3, and 2). Interestingly, we also observed that with increased biliary pressure, some of the blebbing intrusions did not retract back but pinched off from the canalicular surface as larger vesicles (2-8 µm) (Figure 4C,4E and 3). Henceforth, we refer to these vesicles as bile canaliculi-derived vesicles (BCV). Together, these experiments indicate that IB and BCV are cytomechanical responses to increased biliary pressure.

**Figure 3:**
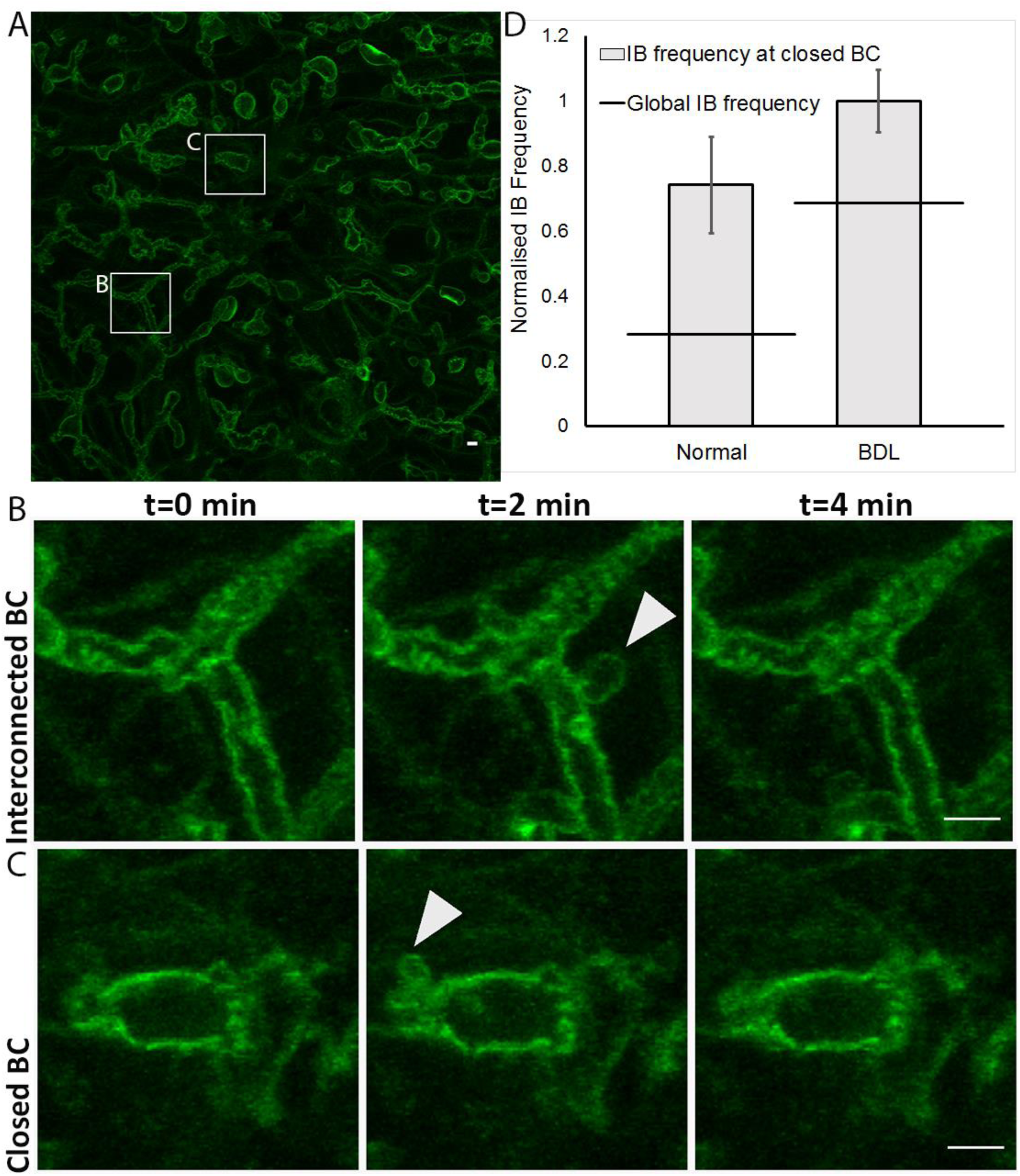
Inward blebs are no longer confined to closed or isolated bile canaliculi after bile duct ligation but also found along interconnected bile canaliculi. (A) shows a Z-projection of confocal slices taken of the liver in bile duct-ligated GFP-LifeAct mice (a representative slice in the stack is shown in Figure 1F). Inward blebs were observed along long and interconnected bile canaliculi (highlighted by square labeled B) and at the terminal ends or along short and isolated bile canaliculi (highlighted by square labeled C). A) and (C) show time lapse images (at 2 min intervals) of the regions highlighted by the squares in (A). Both interconnected bile canaliculi (B) and closed bile canaliculi (C) undergo inward blebbing. Supporting Figure 4 illustrate how interconnected and closed bile canaliculi in bile duct-ligated mice were determined. White arrowheads in (B) and (C) indicate the formation of inward blebs. All scale bars in (A), (B) and (C) are 5 µm. (D) shows quantification of inward blebbing frequency in the bile canaliculi of normal and bile duct-ligated mice.

**Figure 4:**
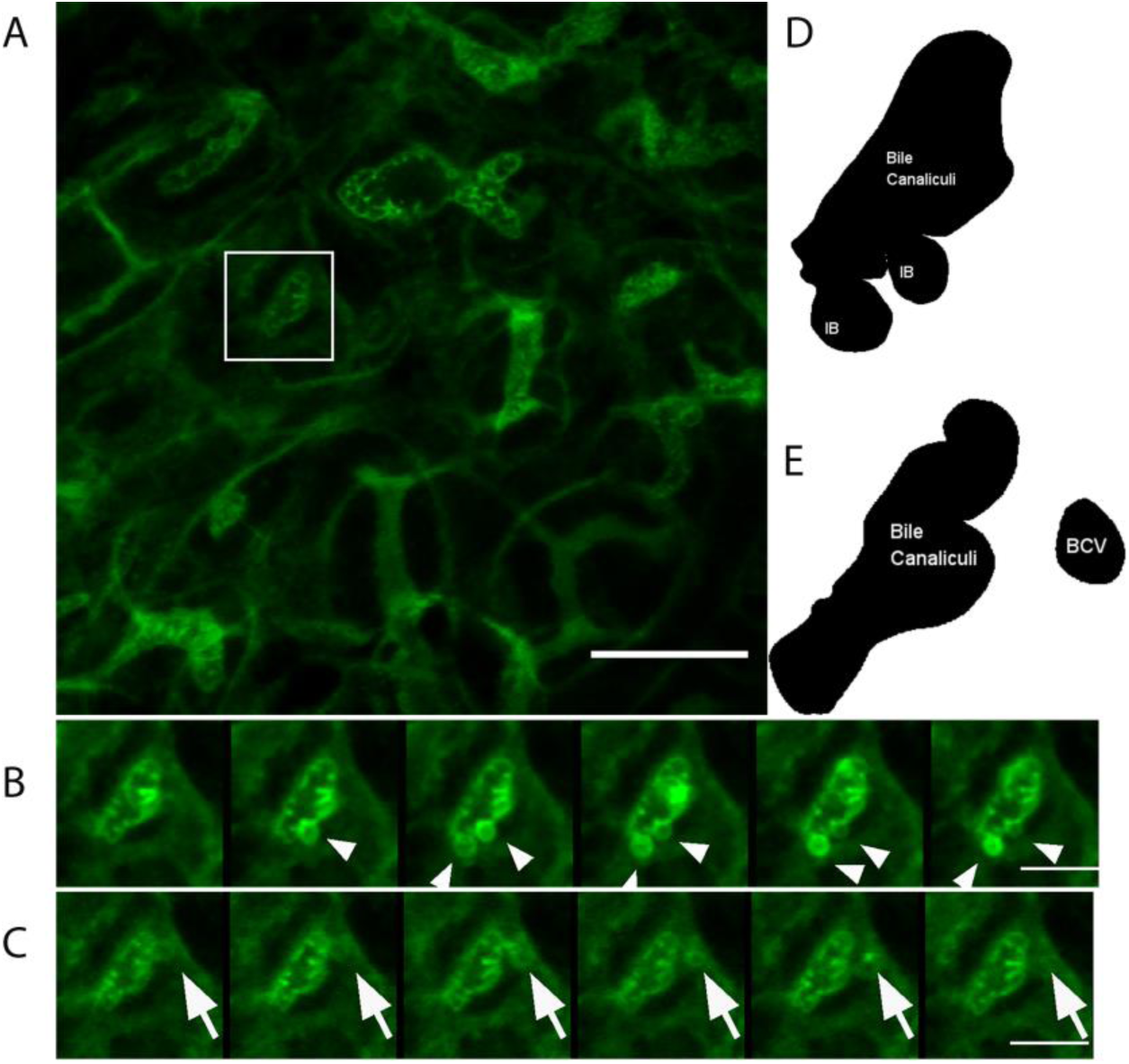
Inward blebs bud off as vesicles into the hepatocyte cytoplasm in bile duct-ligated mice. In bile duct-ligated mice, inward blebs (IB) were often observed to bud off as bile canaliculi-derived vesicles (BCV). Time lapse images (at 2 min intervals) of the region highlighted by the square in (A) is shown in (B) and (C). (B) shows the formation of inward blebs (white arrowhead) in a bile canaliculus and (C) shows the pinching off of an inward bleb (white arrow) as a BCV. (D) Binary schematic of blebbing bile canaliculus in (B). (E) Binary schematic of BCV that has just detached away from the bile canalicular surface in (C). Scale bar in (A) = 25 µm, (B) and (C) = 10 µm.

### IB and BCV formation are triggered at maximum BC area

Given that the expansion of classical extruding blebs is closely associated with cortical actomyosin contraction,[30] we hypothesized that IB and BCV similarly result from a dynamic interplay between ICP and pericanalicular actomyosin cortex (PAC) contractility. To facilitate experimental manipulation and the observation of subcellular dynamics by high-resolution microscopy, we focused our subsequent investigations of IB and BCV in the collagen sandwich culture system (Figure 5A). It is well established that primary rat hepatocytes cultured in this configuration form functional BC that enable the study of bile canalicular dynamics as BC exhibit periodic contraction cycle similar to contraction pattern observed *in vivo* (Supporting Figure 5, s 4 and 5).[3, 21, 22, 31, 32] We first characterized the temporal relationship between IB and BCV formation with specific phase(s) of the bile canalicular contraction cycle. Imaging the GFP fusion of the f-actin probe (F-tractin) that was transfected into rat hepatocytes, we observed that during a typical BC contraction cycle, the BC area increased during the expansion phase without significant changes in the intensity of the PAC until a maximum area was reached (Figure 5E). Following this point of inflection, we then observed the accumulation of f-actin in the PAC and the concurrent appearance of IB and BCV. Notably, on closer examination of the PAC, we noticed that the inward blebbing events were initiated by local ruptures in the PAC, which resulted in herniations in the pericanalicular membrane (Figure 5C and 5D). These results suggest that the PAC serves to provide a counter-balancing force against increasing ICP. Above a certain threshold of ICP, the PAC begins to rupture locally, resulting in the formation of IB (Figure 5D-F and 5). Thus, our data demonstrates how, in the low-pressure regime, the PAC may play a relatively passive mechanical role in supporting BC membrane integrity, while beyond a pressure threshold, the PAC and the BC membrane switch to a highly dynamic actively remodeling mode to adapt to greater mechanical loads.

**Figure 5:**
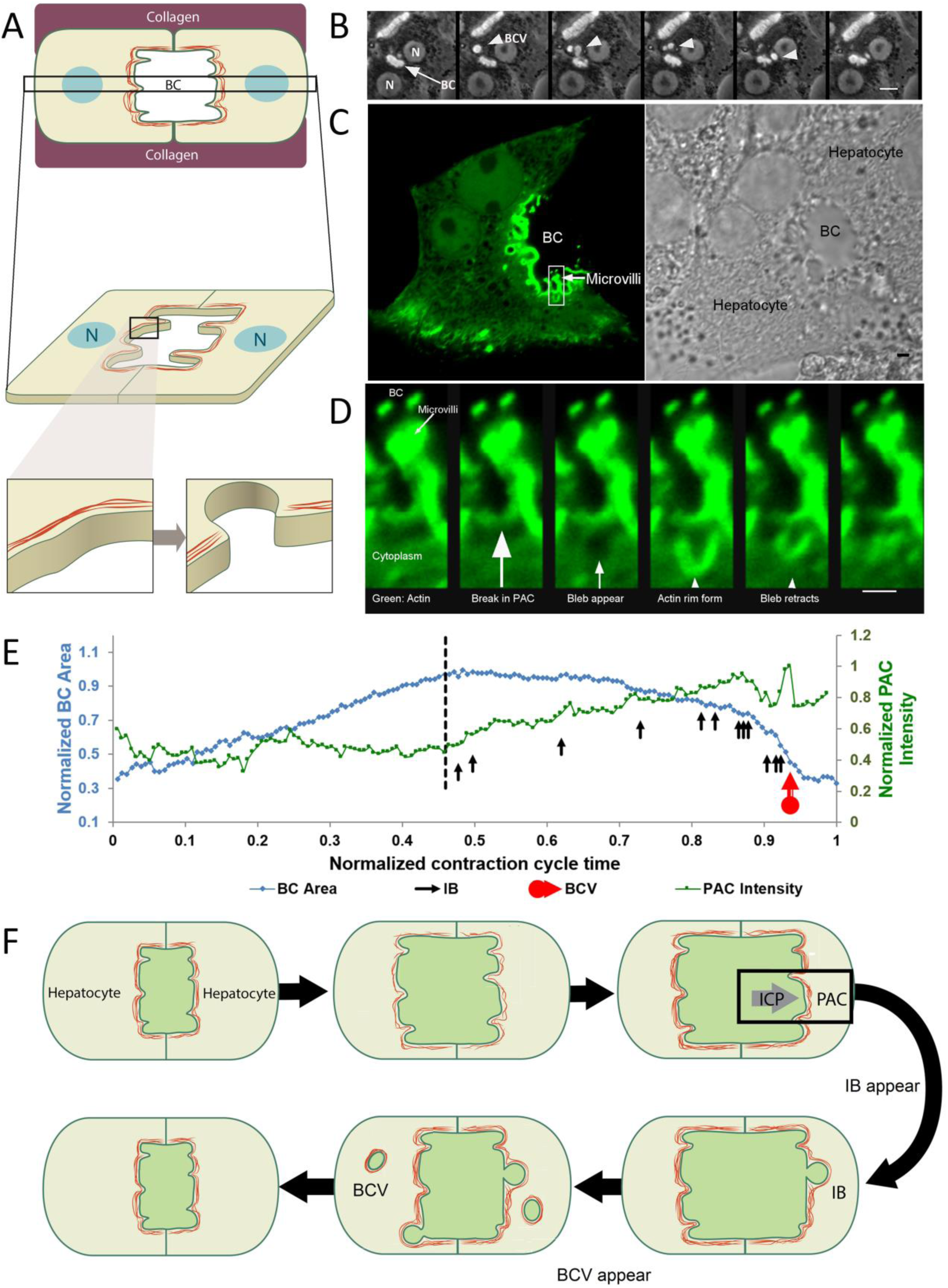
Inward bleb and BCV formation occurs following a break in the pericanalicular actin cortex, as observed in collagen sandwich culture. (A) is a schematic of the collagen sandwich culture system employed to monitor the inward blebbing process in real-time. (B) shows a time lapse (at 7.5 min interval) progression of BCV formation (white arrowhead) from a bile canaliculus (white arrow) in hepatocytes cultured in the collagen sandwich culture system. The hepatocyte nucleus is represented by ‘N’ and the bile canaliculus as ‘BC’. (C) shows a GFP-f-Tractin-transfected hepatocyte (marker for actin, shown in green) forming a bile canaliculus with another non-transfected hepatocyte (as shown in adjacent bright field image). (D) shows a high magnification time lapse (at 26s interval) of the region highlighted by the square in (C). A local rupture in the PAC is observed before a membrane intrusion into the hepatocyte cytoplasm occurs. Following which, an actin rim forms along the membrane bleb and the bleb retracts back towards the bile canalicular space. Scale bar in (B) = 10 µm, (C) = 2 µm and in (D) = 1 µm. (E) Changes in the area of a bile canaliculus and corresponding PAC intensity over time. PAC starts to increase in intensity only after a maximum bile canalicular area is reached (dashed black line). Like-wise, inward bleb (black arrows) and BCV formation (red arrow) were observed to occur only at maximum bile canalicular area and when PAC intensity started to increase. (F) Schematic of events that lead to formation of IB and BCV.

### Altered PAC dynamics influence IB and BCV formation

Given the observation that IB and BCV formation are positively correlated with elevated biliary pressure *in vivo*, we next sought to investigate whether the frequency of IB and BCV formation is dependent on ICP.[33] We experimentally increased ICP by stimulating bile acid secretion using the bile acid, ursodeoxycholic acid (UDCA). Corroborating what was observed *in vivo* with bile duct ligation (Figure 3 and 4), we observed an increase in the frequency of both IB (4-fold increase at 100 µM UDCA) and BCV (9-fold increase at 100 µM UDCA) with UDCA treatment (Figure 6A and B), suggesting that at higher pressure, more IB are unable to retract. These IB then pinch off the canalicular membrane as BCV into the hepatocyte cytoplasm. We next investigated the contribution of the actin cortex to IB and BCV formation by using pharmacological disruptors of the actin cortex. In the presence of cytochalasin D, an inhibitor of actin polymerization,[34] we observed that the frequency of IB formation decreased monotonically with increasing concentrations of cytochalasin D (Figure 6C). However, the frequency of BCV formation exhibits a biphasic response, increasing until 0.2 µM and then decreasing above 0.2 µM (Figure 6D and 6 and 7). Since actin polymerization along the naked IB membrane was observed to occur prior to bleb retraction (Figure 5D and 7A), the inhibition of actin polymerization by cytochalasin D likely impaired the process of IB retraction and therefore, facilitated the formation of BCV. In contrast, at high concentrations of cytochalasin D, the general weakening of the entire PAC likely inhibited any herniation of the bile canalicular membrane (7). In support of this, similar trends were observed when blebbistatin was used to inhibit myosin activity (Figure 6E and F, 8 and 9).[34]

**Figure 6:**
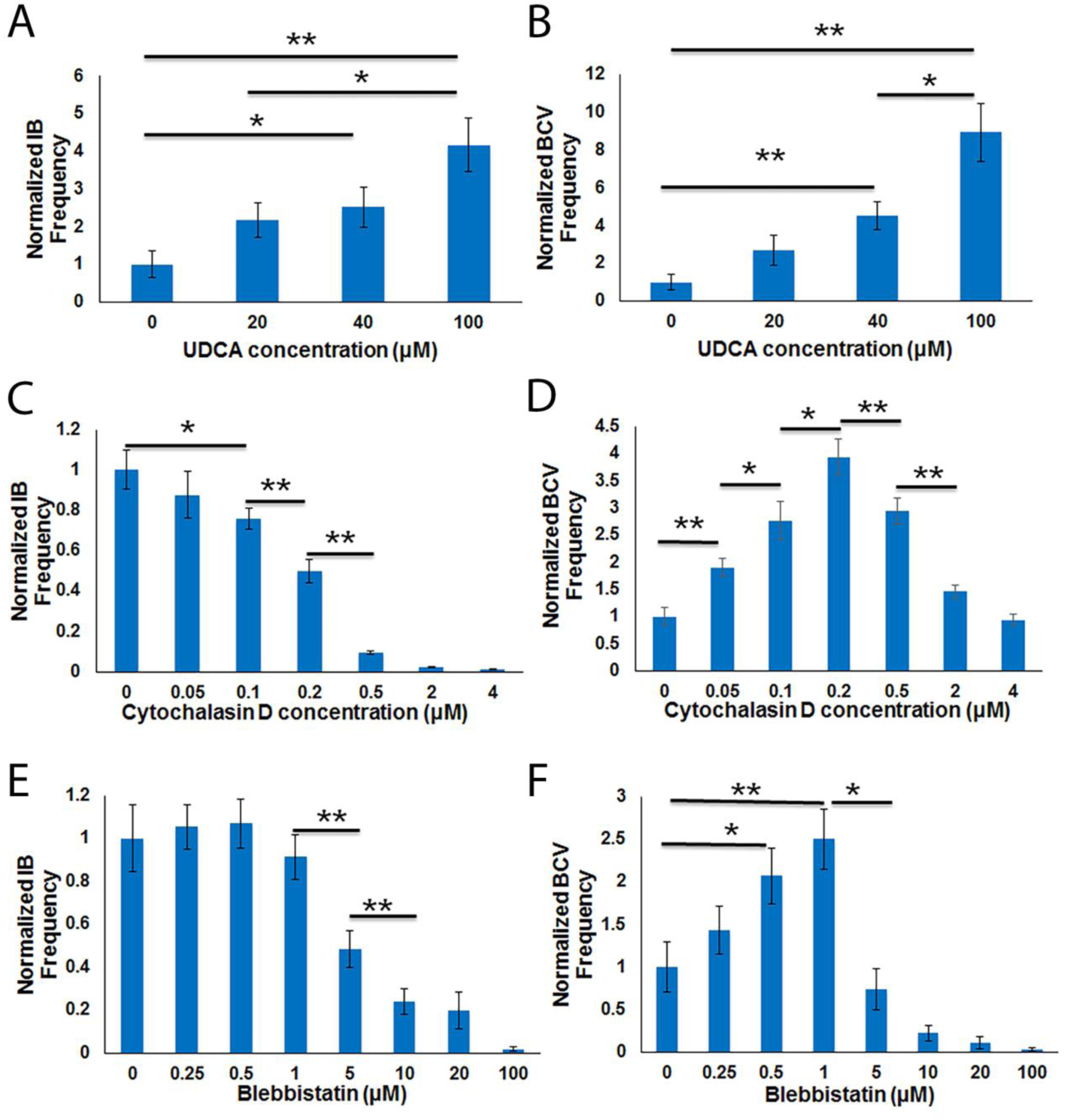
Canalicular pressure increase and perturbations to the pericanalicular actin cortex affect formation of inward blebs (IB) and bile canaliculi-derived vesicles (BCV). Frequency of both IB (A) and BCV (B) increase with increasing canalicular pressure. Canalicular pressure was increased by using UDCA to stimulate bile acid secretion. In the presence of increasing concentrations of cytochalasin D, an inhibitor of actin polymerization, IB formation decreases (C) while that of BCV increases then decreases (D). A similar trend is seen in the presence of blebbistatin, an inhibitor of myosin activity as shown in (E) and (F). *p* < 0.01. \**p* < 0.05, \*\**p* < 0.01. N=40 for all conditions analyzed.

**Figure 7:**
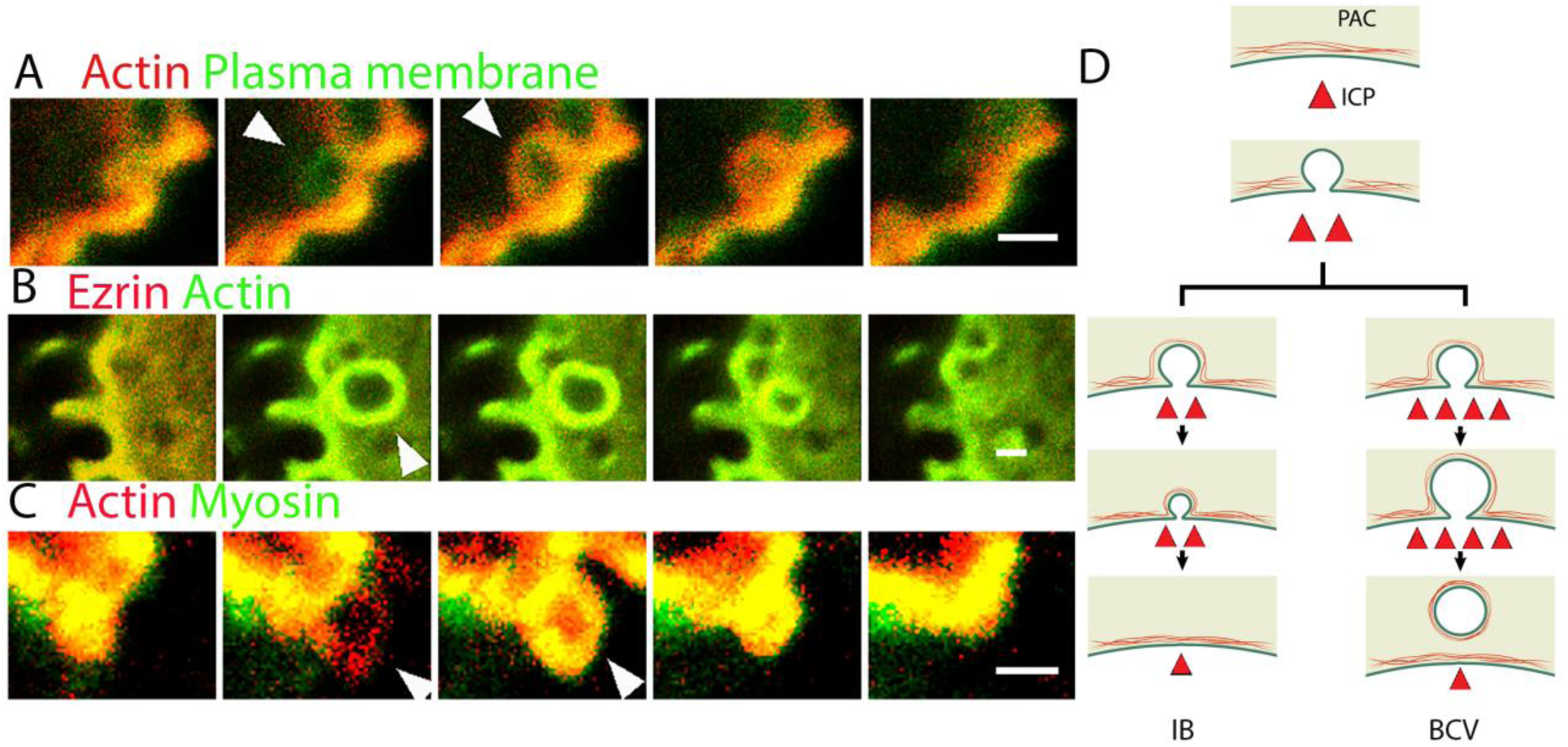
Molecular mechanism underlying inward bleb formation is similar to classical blebbing. Time lapse images of inward bleb formation along the bile canalicular membrane. (A) Following a local rupture in the pericanalicular actin cortex (as shown in Figure 2B), an inward bleb is initiated along the canalicular membrane (green) and expands into the hepatocyte cytoplasm (white arrowhead in (A)). Following which, actin (red) is recruited. The yellow-orange overlay is a result of actin and plasma membrane co-localization. (B) Actin (green) recruitment is accompanied by ezrin (red) (white arrowhead in (B)). The yellow-orange overlay is a result of actin and ezrin co-localization. B) Actin (red) recruitment is followed by myosin (green) recruitment (white arrowhead in (C)). The yellow-orange overlay is a result of actin and myosin co-localization. Stack interval in (A) = 14 s, (B) = 11.5 s and (C) = 42 s. Scale bars = 1 µm. (D) Schematic of pericanalicular actin cortex (PAC) response to increased canalicular pressure (ICP), leading to the formation of either inward blebs or bile canaliculi-derived vesicles.

**Figure 8:**
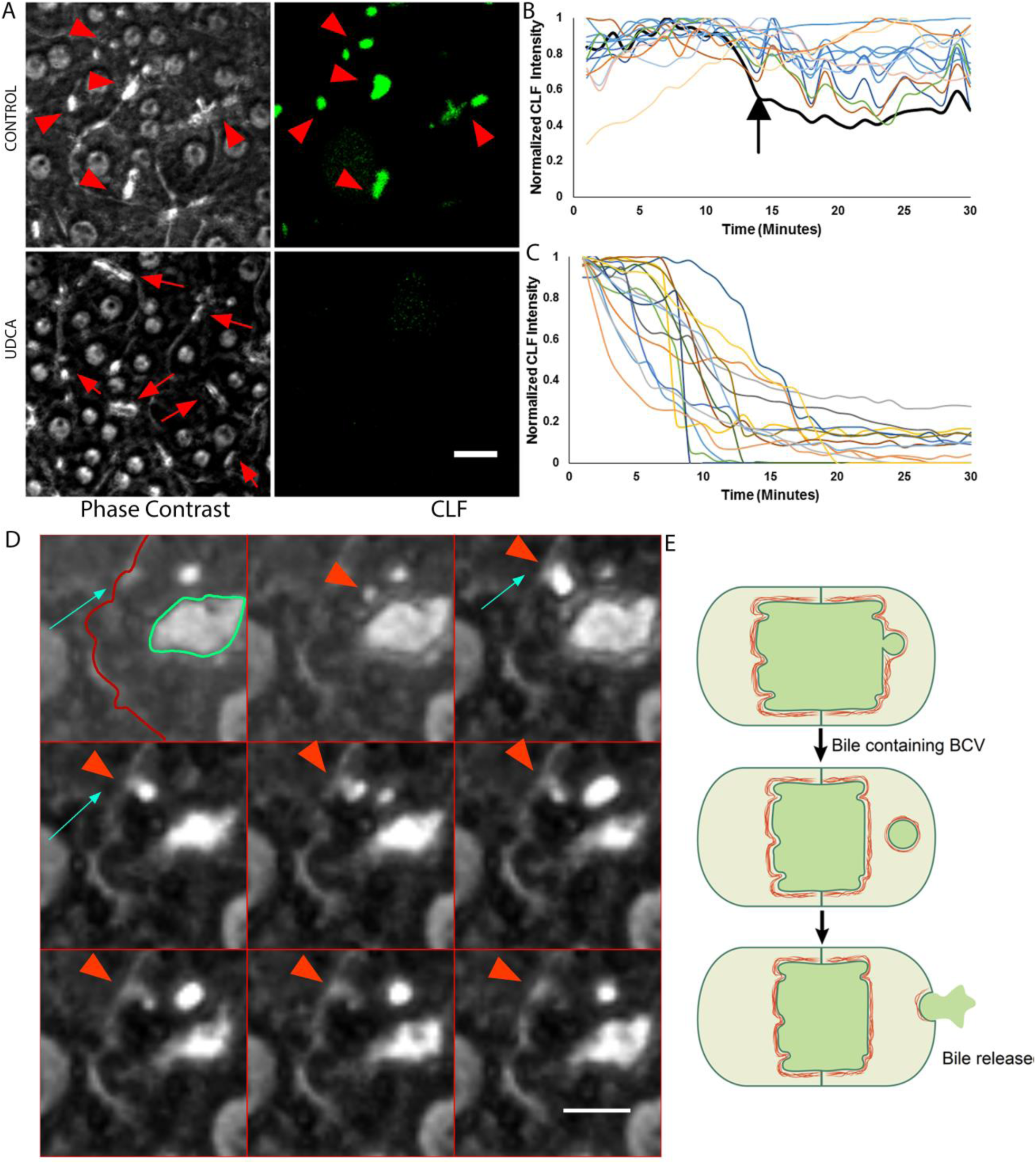
Bile canaliculi-derived vesicles (BCV) enables bile regurgitation during increased canalicular pressure. (A) Using CLF to represent bile acids, CLF was not cleared from the bile canalicular space under normal conditions (arrowhead in (A)) but was rapidly cleared in the presence of UDCA (arrow in (A)). UDCA enables simulation of increased canalicular pressure as it increases bile acid secretion. Scale bar = 20 µm. (B) and (C) show quantification of CLF intensity within bile canaliculi under normal conditions (B) and in the presence of BCV-stimulating UDCA (C). Each curve in (B) and (C) represents CLF intensity in individual bile canaliculi normalized with respect to maximum intensity during 30 min for each bile canaliculi. There were a few bile canaliculi which show a drastic decrease in CLF intensity (black line) under normal conditions (B) but those were bile canaliculi in which BCV formation was observed. 15 individual bile canaliculi were analyzed in (B) and (C). (D) Montage showing release of bile canaliculi-derived vesicle (red arrowhead) at cell edge (red line marked by blue arrow in first frame). As can be seen, BCV travels toward the cell edge and merges with the cell membrane. Scale bar = 10 µm. (E) Schematic of a proposed model of BCV-facilitated bile regurgitation.

### Hepatocytes retract IB by recruiting and contracting actomyosin

Classical extruding blebs commonly occur during apoptosis, cell spreading, and cytokinesis, typically following a three-phase life cycle: bleb initiation, expansion and, cortex repolymerization and retraction.[23, 35–38] Extruding blebs are thus initiated either by a local detachment of the cortex from the cell membrane, or from a local rupture in the cortex. [24] This is followed by cytoplasmic pressure-driven bleb expansion, resulting in a local naked membrane unsupported by an underlying cortex. In the third phase, actin, myosin and other cortical proteins are recruited to the bleb membrane, thus reforming the membrane-anchored actin cortex that enables bleb retraction. [24, 34] To identify the molecular mechanism underlying IB retraction, we expressed fluorescent protein reporters for actin, ezrin (a cortical protein that links membrane to the actin cytoskeleton), plasma membrane and myosin IIA in rat hepatocytes and analyzed their live-cell dynamics using confocal microscopy. Samples were imaged at a distance of at least 2 µm from both the top and bottom of the cell surface to capture events occurring within the cell near the bile canalicular membrane (Figure 5A). We found that subsequent to a rupture in the PAC (Figure 5D and 7A, Supporting Figure 6A and 10), we observed a rapid inward expansion of the naked pericanalicular membrane - devoid of actin and myosin II - intruding into the cytoplasm. Subsequently, actin and ezrin (Figure 7B, Supporting Figure 6B and 11) were recruited to the cytoplasmic surface of the bleb, followed by myosin IIA (Figure 7C, Supporting Figure 6C and 12). These results indicate that IB retraction is driven by the recruitment and contraction of the PAC, analogous to the classical blebs that occur during apoptosis and cytokinesis, the recruitment and contraction of the PAC drives IB retraction. Notably, with further increase in ICP, such as in the presence of high concentrations of UDCA, we observed the detachment of IB from the canalicular membrane as BCV. This likely occurs when the newly formed IB actomyosin cortex is unable to generate sufficient force to retract back into the canalicular space (Figure 7D and 5).

### BCV formation enables bile regurgitation during increased ICP

Based on the observation that IB detach from the canalicular surface as BCV at high ICP, we asked if BCV formation serves a physiological role in biliary homeostasis. Interestingly, we discovered that hepatocytes under UDCA treatment (UDCA increases ICP and hence, stimulates BCV formation) exhibited a significant increase in bile acid clearance from the canalicular lumen as compared to the controls, as assayed using the fluorescent bile acid, CLF (Figure 8A and 13 and 15). Similar results were observed when cytochalasin D was used to stimulate BCV formation (Supporting Figure 7A). To characterize bile clearance dynamics, we then performed live cell fluorescence microscopy, and monitored the levels of CLF in individual BC. Remarkably, in the presence of UDCA, we observed that CLF was nearly completely removed from the BC within 20 min (Figure 8C) via CLF-containing BCV. In contrast, CLF largely remained in the BC in the controls (Figure 8B). In the few cases in the control group where we did observe CLF clearance, these coincided with the rare formation of BCV (Figure 8B, black line marked by arrow; and 14).

As a significantly greater amount of CLF was detected in the surrounding culture medium in the presence of UDCA or cytochalasin D (Supporting Figure 7B), we next asked whether the BCV plays a role in transporting bile acids from the bile canalicular space to the cell exterior as a means of bile regurgitation. First, to eliminate the possibility that BCV contents are not released intracellularly, we monitored the net intensity of entire cells starting from the time when CLF-containing BCV detaches from the bile canalicular surface and enters the cytoplasm, to when BCV disappears. We found that the net intensity of the entire cell increases when the BCV enters the cell, but decreases when the BCV disappears (Supporting Figure 8). If CLF was released within the cell, we would not have observed a reduction in net intensity but rather, unchanging net intensity after the BCV enters the cell. Hence, this indicates that CLF is released to the cell exterior and likely BCV-transported. Indeed, in monitoring individual BCV over time using live-imaging, BCV were observed to translocate from the bile canalicular surface to the cell periphery and appears to come into very close proximity with the plasma membrane (Figure 8D). Lastly, the role of BCV as transporters of bile acid cargo was verified *in vivo* using transmission electron microscopy performed on BDL mouse liver. We found large BCV-like structures pinching off from the bile canalicular surface and also observed similar structures fusing with the sinusoidal membrane (Supporting Figure 9). To eliminate the possibility that BCV were apoptotic bodies, we incubated hepatocytes with NucView™ 530 Caspase-3 substrate and confirmed that BCV were not apoptotic bodies (Supporting Figure 10). Taken together, these results are indicative of the physiological role of BCV formation as a major cellular mechanism for bile regurgitation, serving as a homeostatic ‘relief valve’ in response to increased canalicular pressure (Figure 8E).

## Discussion

The results of our study demonstrate that beyond global BC motility at the tissue level, local cellular events in the form of IB and BCV occur along the bile canalicular surface at elevated canalicular pressures (Figures 1–4), challenging our existing understanding of BC dynamics. The comprehensive tracking of the spatio-temporal response of the canalicular surface to increased pressure enabled us to elucidate in details the major phases of this homeostatic response. In the first phase, before a certain threshold in canalicular pressure is reached, the bile canalicular surface increases in area without any accompanying increase in the volume of the PAC. In the second phase, once the BC surface reaches a maximum area, the PAC starts to thicken to resist the increasing canalicular pressure. During this phase, local contractions in the PAC result in ruptures that cause herniations in the bile canalicular membrane. Under normal conditions, these IB retract back into the bile canalicular space. However, when the canalicular pressure is abnormally high, as in the case of obstructive cholestasis, these IB are unable to retract back into the bile canalicular space and pinch off as large BCV.

The phenomenon of inward blebbing as a cellular response to high external pressure was only very recently reported by Gebala and colleagues, where it was found that blood flow drives lumen expansion during sprouting angiogenesis *in vivo*, by inducing such blebs along the apical membrane of endothelial cells.[26] Similarly, in this study, we establish that increased luminal pressure induces IB along the canalicular membrane, which likewise undergo actomyosin-mediated retraction. However, contrary to the findings in endothelial cells, we observed that the hepatocyte IB do not retract back when the canalicular pressure is sufficiently high enough to cause these blebs to pinch off as BCV. The presence of vesicles or vacuoles in the hepatocyte cytoplasm as a distinct morphological outcome of obstructive cholestasis is not a new observation.[12, 39] Indeed, previous studies have described the presence of vesicles that were functionally associated with transcytosis,[40, 41] movement of transporters to or from the canalicular membrane [39] or endocytosis.[40, 42] Notably, the BCV we observed in this study are large (approximately 5 µm in diameter), unlike previously reported vesicles (less than 500 nm in diameter). Additionally, these studies typically deduced the origin and function of the vesicles based on static electron micrographs. Through these ‘snapshots’, it has been suggested that some of these vesicles are derived from the bile canalicular surface and function as vehicles for bile regurgitation from the bile canalicular surface to the sinusoidal surface when canalicular pressure is increased. However, despite these studies, whether these vesicles indeed originate from the bile canalicular surface and serve to regurgitate bile remained unclear. In this study, we demonstrate that BCV indeed originate from the bile canalicular surface and function to regurgitate bile out of the hepatocyte (Figure 8), therefore confirming previous correlative studies [11, 43] and establishing for the first time, this vesicle-mediated transcellular pathway as an important mechanism of bile regurgitation. Notably, studies are ongoing in our laboratory to elucidate the exact mechanism of bile release when the BCV is at the basolateral surface; BCV can either release bile contents to the cell exterior by fusing with the cell membrane or, do so via the ‘Kiss and Run” model [44] where it momentarily comes into contact with the cell membrane to release bile but the membrane is subsequently recovered. As transporter proteins are rarely found on the basolateral membrane in cholestatic conditions [45], the ‘Kiss and Run’ model is the likely mechanism that regulates this process.

In seeking to understand the molecular machinery driving the formation of IB and BCV, we analyzed the global BC contraction cycle in detail and observed that the accumulation of PAC and emergence of IB only began when the maximum BC area was reached during the expansion phase. Tantalizingly, these results are suggestive of the participation of mechanosensors and mechanotransducers as limiting switches; these switches may be involved in triggering homeostatic cellular responses to limit any further increase in canalicular pressure that might compromise cell-cell contact and tissue integrity. Indeed, actin polymerization in response to increased intra-vascular pressure was previously described in vascular smooth muscle cells in response to increased intravascular pressure.[46] As adherens junctions are coupled to the actin cytoskeleton to enable the transmission of actomyosin forces between adjacent cells in epithelial cell sheets, we surmise that the adherens junctions could serve as mechanosensing sites during this process.[47] Indeed, Yonemura and colleagues showed that α-catenin functions as a tension transducer, undergoing force-dependent conformational changes to recruit the actin-binding protein vinculin.[48] The subsequent increase in the association with actin then strengthens the adherens junction to withstand tensional forces between cells. In the context of this study, we postulate that increased canalicular pressure exerts tensional forces on cell-cell junctions between adjacent hepatocytes, potentially triggering similar mechanisms of mechanotransduction that lead to a strengthened PAC. The precise molecular events that transduce such pressure-induced tensional forces to actin polymerization reported in this study are now being actively investigated in our laboratory.

To counteract an increasing canalicular pressure that a strengthened PAC is unable to suppress, the PAC undergoes local remodeling to accommodate the formation of BCV to regurgitate bile. Above a certain pathological pressure threshold, local herniations in the bile canalicular membrane (IB) cannot be repaired and thus pinch off as BCV. Using agents that either disrupt actin polymerization or myosin activity (Figure 6), we demonstrated that the frequency of BCV formation can be modulated, suggesting that the PAC could be a potential therapeutic target to control BCV formation and hence, bile regurgitation. As BCV is typically observed within 1-2 h after bile duct ligation, it is likely that this process represents an early homeostatic response of a biophysical nature that occurs before the onset of other homeostatic mechanisms, such as changes in transporter activity [49]. Also, as this process appears to be mainly physical in nature, it is likely that BCV unselectively transports other important biliary constituents (including bilirubin and biliary lipids) in addition to bile acids during increased biliary pressure. As an example, it has been previously reported that lipoprotein X appears in the systemic circulation very early in cholestasis, the mechanism through which this occurs has yet been determined. Studies are ongoing to investigate the relevance of BCV-mediated regurgitation in the clearance of biliary constituents other than bile acids, such as lipoprotein X [50]. Lastly, while the removal of bile acids from hepatocytes by BCV may minimize hepatocyte injury and reduce canalicular pressure, the accumulation of bile acids in the systemic circulation via bile regurgitation may contribute to endothelial cell injury in the kidneys and lungs. [51] Further studies are needed to understand how bile regurgitation should be managed to strike a balance between toxicity to the liver versus damage to the other organs in the body. Additionally, how this process of bile regurgitation contributes to the pathogenesis of ductular proliferation, hepatocyte injury and portal fibrosis, which are hallmarks of cholestatic injury, remains to be elucidated. Understanding this process will potentially yield novel treatment options for patients with primary biliary cirrhosis and primary sclerosing cholangitis.

In conclusion, the present study demonstrates how the bile canalicular network responds at the subcellular level to increased canalicular pressure during obstructive cholestasis by strengthening the PAC and inducing the formation of BCV that regurgitate bile. Our results underscore the significance of the PAC in contributing to the described homeostatic responses that function to alleviate obstructive cholestasis. By identifying the exact sequential events leading to the formation of bile-regurgitative BCV, our results suggest potential therapeutic targets that could provide novel strategies for the management of bile regurgitation in patients with obstructive cholestasis.

## Acknowledgements

The LifeAct-GFP mouse model used in this study was a kind gift from Dr. Roland Wedlich-Söldner (University of Munster, Germany). All schematics in the paper were designed by Wong Chun Xi, Illustrator, Science Communications Facility, Mechanobiology Institute, National University of Singapore. We also thank the Electron Microscopy Unit at the Yong Loo Lin School of Medicine, Confocal Unit at the Yong Loo Lin School of Medicine and the Microscopy Core at the Mechanobiology Institute, National University of Singapore for their help with electron and confocal microscopy

